# Genome dynamics across the radiation of a mega-diverse genus

**DOI:** 10.1101/2025.09.17.676772

**Authors:** L Campos-Dominguez, TE Kongsted, L Cure, M Downie, A Martinez-Martinez, C Fan, L-N Dong, Y-H Tseng, A-Q Hu, K-F Chung, J Pellicer, IJ Leitch, A Bombarely, AD Twyford, CA. Kidner

## Abstract

Understanding the drivers of species diversity and rapid radiations is a major goal in evolutionary biology. *Begonia* is one of the most species-rich angiosperm genera with 2,164 species currently identified. This genus exhibits considerable variation in chromosome number and a wide range of genome sizes, allowing us to associate genome dynamics with divergence and speciation at a range of temporal scales. We investigate all main radiations within the Begoniaceae family using five previously published *Begonia* genomes and seven new genome assemblies. We show that *Begonia* species show more complex, repetitive and dynamic genomes overall than their close relative, the monotypic *Hillebrandia sandwicensis*. We identify families of repetitive elements that have recently expanded in species from two different highly speciose Southeast Asian sections and two large Neotropical radiations. Detailed characterisation of genomes from species belonging to two parallel radiations, one in Southeast Asia (*Begonia* section *Coelocentrum*) and the other in the Neotropics (*Begonia* section *Gireoudia*), revealed recent expansion in LTR retrotransposons (LTR-RTs) and satellite DNA, in contrast to more species-poor closely related clades. We further investigate variation in repetitive elements within species, finding that accessions from a population of the widespread *Begonia heracleifolia* with unusually large genomes show a markedly higher satellite repeat and Ty3/Gypsy LTR-RT content associated with the expansion of a few abundant repeat lineages. We find that accessions derived from this population show lower seed viability in crosses with other conspecific populations, and thus identify a direct link between expansions of repetitive DNA and the process of genetic isolation. These results show how genome dynamics may promote speciation in one of the most diverse flowering plant genera.

## Introduction

One of the most distinctive aspects of biological diversity is its uneven distribution across lineages: some groups contain thousands of species, whereas closely related sister lineages may have only a few (Donoghue and Sanderson, 2015). Angiosperms have long been singled out as an example of unusually rich taxonomic diversity emerging in a relatively brief period of time (Heywood, 1993). This species richness is also unevenly distributed within the group: a few angiosperm families include thousands of described species, others include only one or two. Despite difficulties in modelling diversification rates (Louca and Pennell, 2020) it is clear that diversification rates of angiosperms are highly variable (Magallon and Sanderson, 2001; Tietje et al., 2022; Zuntini et al., 2024) and the uneven branching patterns of most angiosperm phylogenies suggests that the drivers of diversification act differently across the plant tree of life. One of the major challenges in the field of evolutionary biology is describing and analysing the factors driving speciation, however doing so will elucidate the processes underlying the remarkable differences in species diversification between angiosperm groups.

The most commonly considered factors contributing to species radiations are environmental and ecological (Tao et al., 2021). Nevertheless, intrinsic biological factors have also been identified, which influence the rate of speciation (Marques et al., 2019; Escudero and Wendel., 2020; Heslop-Harrison et al., 2023). Leitch and Leitch (2012) proposed that variation and interaction between extrinsic (ecological, environmental) and intrinsic (changes at the genome level) factors will affect genome diversification and evolution, and therefore evolutionary rates. Fundamentally, all processes contributing to adaptation, diversification, and speciation are determined by genomic variation, which can arise through intrinsic, small-scale events (e.g., point mutations, recombination errors) or through more complex evolutionary events such as polyploidization, hybridization or Transposable Elements (TE) bursts. This variation is the source material of evolution, which through either adaptive or non-adaptive processes can generate genetic incompatibilities, leading to population divergence and eventually speciation (Mérot et al., 2020; Bowles et al., 2020).

Repetitive DNA, such as TEs and satellite DNA (satDNA) (regions of tandemly repeated short repetitive sequences) are major contributors to genomic variation between lineages. In fact, in angiosperms, the huge variation in genome size (Pellicer et al., 2018) is largely accounted for by variation in the length and number of repetitive elements (Maumus and Quesneville, 2014). TE activity has long been known to cause severe changes in genomic architecture (McClintock, 1953; Kidwell and Lisch, 1997). These elements attain mobility in the genome through processes in which they are copied and transposed, keeping both the old and new positions (class II elements or retrotransposons) or are cut out and transposed into a new position (class I TE or DNA elements). Their structure includes coding regions that make these processes autonomous. Recently, TEs have been proposed to play a more important role than polyploidy in plant genome size variation and evolution (Ibarra-Laclette et al., 2013; Bennetzen and Wang, 2014; Catlin and Josephs, 2022). Although TEs and satDNA present structural differences, sequence similarity has led to the hypothesis that satDNA can be derived from TEs (Mestrovic et al., 2015), making regulation of TE activity a key control point in genome structure.

The recent increased affordability of genome sequencing has enabled research on TE content and genome structure in an increasingly wide range of angiosperm lineages (e.g. Novak et al. (2014); Beule et al. (2015); Zhang et al. (2015); Niu et al. (2019); Belyayev et al. (2020)). The roles of TEs in evolution have now been studied in many different angiosperm genera, including *Nicotiana* (Parisod et al., 2012), *Solanum* (Dominguez et al., 2020), *Aegillops* (Senerchia et al., 2013, 2015), *Biscutella* (Bardil et al., 2015), *Oryza* (Zhang Gao, 2017; Akakpo et al., 2020), *Helianthus* (Mascagni et al., 2017), *Cucumis* (Morata et al., 2018), *Arabidopsis* (Quadrana et al., 2019) and *Citrus* (Meng et al., 2020). However, most of these studies focus on the evolution of crop species, small taxa or on specific TE families. Although they are widely considered one of the main sources of genetic variation in plants, genome-wide studies on the role of the repeatome in the evolution of large genera are still rare.

New genus-wide studies on the factors driving large angiosperm radiations, particularly those making use of the flood of genome-level data, can give us new perspectives on the evolutionary drivers of mega-diversity (Seehausen et al., 2014). For this purpose, the genus *Begonia* L. is proving to be a particularly suitable model. This pan-tropically distributed genus has very similar (both geographical and taxonomical) patterns of species richness to angiosperms overall (data from Hughes et al. (2015)), and is among the five largest genera of flowering plants, with many new species being described every year (2,164 described species, as stated in the Begonia Resource Centre Database from Hughes et al. (2015) in Oct 2024). In terms of their biogeographic history, two major recent radiations in the Neotropics and in Asia respectively are derived from an early diverging African *Begonia* lineage (Goodall-Copestake et al., 2010). *Begonia* species are known for their wide phenotypic diversity, particularly in leaf shape (Neale, 2005; Ali, 2013). They grow in a wide range of habitats with varying light and humidity levels, though are most common in the high humidity, moderate temperatures of montane rainforests (Burt-Utley, 1985).

The mega-diversity of *Begonia* contrasts with its only sister genus in the family Begoniaceae, *Hillebrandia* Oliv. (Hughes, 2002), which has only a single species (*H. sandwicensis*), restricted to the Hawaiian Islands (Clement et al., 2004). Within *Begonia*, the early divergent clades from mainland Africa are relatively species-poor. In the Neotropics, Asia and Madagascar, the genus is exceptionally rich and exhibits considerably higher speciation rates compared to the African sister clades (Moonlight et al. (2015, 2018)). Dewitte et al. (2009) and Campos-Dominguez et al. (2022) documented genome size and chromosome number variation in *Begonia*, suggesting the dynamic nature of the *Begonia* genome may contribute to speciation in a non-adaptive speciation model (Rundell and Price 2009). In this study we explored the dynamics of genome repeats across *Begonia* evolutionary radiations at different timescales. Our overall aim is to test the role of genome dynamics in the origins of diversity within the Begoniaceae, by both documenting the repeated outcomes of TE variation in diverse species radiations, and by testing whether similar variation manifests within species and may drive divergence at the population scale.

For this purpose, we sequenced and assembled the genomes of seven new species, including high-quality long-read based assemblies for *H. sandwicensis* from the *Begonia* sister genus *Hillebrandia*, and two Neotropical *Begonia* species. We further investigated the genomic repeat dynamics across section-level radiations within Asian and Neotropical clades, as well as within the species *Begonia heracleifolia*. We hypothesize that TE activity and changes in the repeatome are associated with population divergence and speciation, and report TE content variation associated with diversification between populations of a widespread species (*B. heracleifolia*), as well as expansions of specific repeat families in diverse *Begonia* clades.

## Materials and methods

### Plant material

A full list of all species’ accession numbers and collection of origin can be found in supplementary table 1. Flower and leaf buds of the five *Begonia* species used for whole genome sequencing and assembly (*B. dregei, B. socotrana, B. bipinnatifida, B. johnstonii* and *B. luxurians*) were collected from the *Begonia* living collections at the Royal Botanic Garden Edinburgh (RBGE) and kept in silica gel or frozen in liquid nitrogen. The *Hillebrandia sandwicensis* silica-dried flower tissue samples and seeds were sent from the National Tropical Botanical Gardens in Hawaii (USA). In the case of the specimens used for low-coverage sequencing (genome skimming), either flash-frozen or silica-dried tissue was used for DNA extractions using the Qiagen plant minikit (catalog number 69104). These were all part of the living collections of the Biodiversity Research Center in Academia Sinica (Taiwan) and the RBGE. *B. oxacana* material was provided by the Royal Botanic Gardens, Kew (RBG Kew).

### C-value estimations of B. heracleifolia populations

We estimated the C-values of 10 individuals from different populations of *B. heracleifolia* originally collected in Mexico and kept growing at the RBGE living collections (Twyford et al., 2014). Three fresh leaves were collected from each individual and analysed independently. Nuclear DNA contents (C-values) were measured at the Jodrell Laboratory at the RBG Kew. Briefly, we followed the two-step protocol described in Pellicer & Leitch (2014), using *Solanum lycopersicum* ‘Stupiké polní rané’ (2C=2.0 pg (Praça-Fontes et al., 2011)) as a reference standard, and the CysStain PI Absolute nuclei extraction buffer (Sysmex, Kobe, Japan). Samples were analysed using a CyFlow SL3 Partec flow cytometer (Sysmex-Partec, Munster, Germany) fitted with a 100 mW green lamp (532 nm solid-state Cobalt Samba laser; Cobolt AB, Solna, Sweden). Three replicates per individual were prepared and run three times each. The resulting flow histograms were analysed using the Partec software for flow cytometry FloMax 2.9 (Sysmex-Partec).

### DNA extraction, library preparation and sequencing

We initially obtained short-read Illumina sequencing data for all samples, then assembled genomes from a subset of species using different long-read sequencing approaches for samples of particular interest for their phylogenetic position (e.g. Neotropical *Begonia* and *Hillebrandia*), or where there were potential challenges caused by high heterozygosity (which would make short-read assemblies harder to resolve). We then obtained data using Pacific Biosciences (Pacbio RSII, GenomeQuebec) sequencing and Oxford Nanopore Technologies (ONT) of *B. conchifolia*, Pacbio HiFi of *B. luxurians* and ONT sequencing of *H. sandwicensis*. Detailed methods and sequencing stats are in supplementary methods.

### Genome assemblies

The *B. conchifolia* genome was assembled using the Pacbio data with Flye 2.4.2 (Kolmogorov et al., 2019) and the resulting assembly was polished and re-scaffolded using Illumina and ONT data, respectively. The Pacbio HiFi reads of *B. luxurians* were assembled using Hifiasm-0.18.5 (Cheng et al., 2021), and haplotype duplicates were removed using purge_dups v1.2.6 (Guan et al., 2020). The genome assembly of *H. sandwicensis* was performed using Soapdenovo r241 (Luo et al., 2012; sup fig 2), re-scaffolded using ONT reads (with Porechop v0.2.4; Wick, 2018 and the SLR software; Luo et al., 2019) and polished (with Medaka v1.7.3; Oxford Nanopore Technologies and polypolish v0.5.0; Wick & Holt, 2022) using the short-read data.

We also generated whole genome short read sequencing data for four additional species to survey genome dynamics over a wider range of taxa *(B. dregei, B. bipinnatifida, B. socotrana*, and *B. johnstonii)*. Best assemblies based on quality metrics (N50, number of contigs/scaffolds, completeness and BUSCO scores) were selected and afterwards cleaned for short and contaminant sequences using blobtoolkit v2.2 (Challis et al., 2020).

### Transposable Element annotation

The Extensive de novo TE Annotator (EDTA v2.1.0; Ou et al., 2019) was used to find and characterise TEs in all the available and newly generated genomes, and annotated elements (TEanno output) were further characterised using TEsorter v1.4.6 (Zhang et al., 2022). Insertion times were obtained from the LTR-retriever output generated by EDTA.

For cluster-based repeat annotation of the low-coverage data, only the short raw reads were used. The *Begonia foliosa* var. *miniata* reads were downloaded from the Short Nucleotide Archive (SRA study code SRP100595; Griessman et al., 2018). For all species, the number of trimmed and clean reads equivalent to 0.2x coverage were randomly sampled for each dataset using seqtk 1.3 (Li, 2012). Reads were then uploaded, pre-processed, clustered and annotated by similarity using the RepeatExplorer2 pipeline, (Novak et al., 2020) implemented in a Galaxy server environment (https://repeatexplorerelixir.cerit-sc.cz). For the samples that showed high satellite DNA content, a second RepeatExplorer2 run with satellite filtering parameters was performed. For each of the species, cluster annotations were then reviewed and manually curated, and all clusters identified as plastid or mitochondrial DNA were removed prior to any downstream analysis.

For the comparative analyses, 440,000 random sequences were sampled for each species, pre-processed and code-named to be pooled into the same dataset. Reads were then clustered and annotated using the satellite filtering and comparative analysis RepeatExplorer2 settings.

### Genetic crosses of B. heracleifolia populations and seed viability tests

Crosses between *B. heracleifolia* accessions were performed as described in Twyford et al. (2014). *Begonia* individuals were crossed by hand-pollinating newly opened female flowers with male flowers, with pollinated flowers bagged to prevent pollen contamination. Mature seed capsules were harvested and viability rates determined by scoring the ratio of full seeds in random samples of 100 seeds per capsule. For the analysis of seed viability, crosses were divided into self-crosses (SELF), outcrosses between accessions of a similar genome sizes (OUT_within), outcrosses between accessions of distinct genome sizes (OUT_between), and crosses done using F1 populations as parents (F).

## Results

### Genomic data of Begoniaceae: seven new genome assemblies

We have obtained seven new genomes from across the Begoniaceae family: six *Begonia* species (three from African clades, two American clades and one Asian clade) as well as *Hillebrandia sandwicensis*, the representative from their monospecific sister genus. These genomes were sequenced using different short-read and long-read technologies and approaches. Figure 1 shows the species sequenced in a phylogenetic context and a table with genome data from the seven new genomes as well as the previously sequenced ones (Li et al., 2022; Griessman et al., 2018). These all belong to different *Begonia* sections (except *B. darthvaderiana* and *B. bipinnatifida*), with a variable number of species in each of these sections. *Begonia loranthoides, B. dregeii, B. socotrana* and *B. johnstonii* belong to small, more depauperate African sections with 31, 12, 2 and 9 species, respectively. *B. conchifolia, B. masoniana, B. darthvaderiana, B. bipinnatifida*, and *B. luxurian*s belong to large *Begonia* sections representing radiations of 110, 93, 434 and 235 species respectively (data from Hughes et al.,2015).

**Figure 1.**
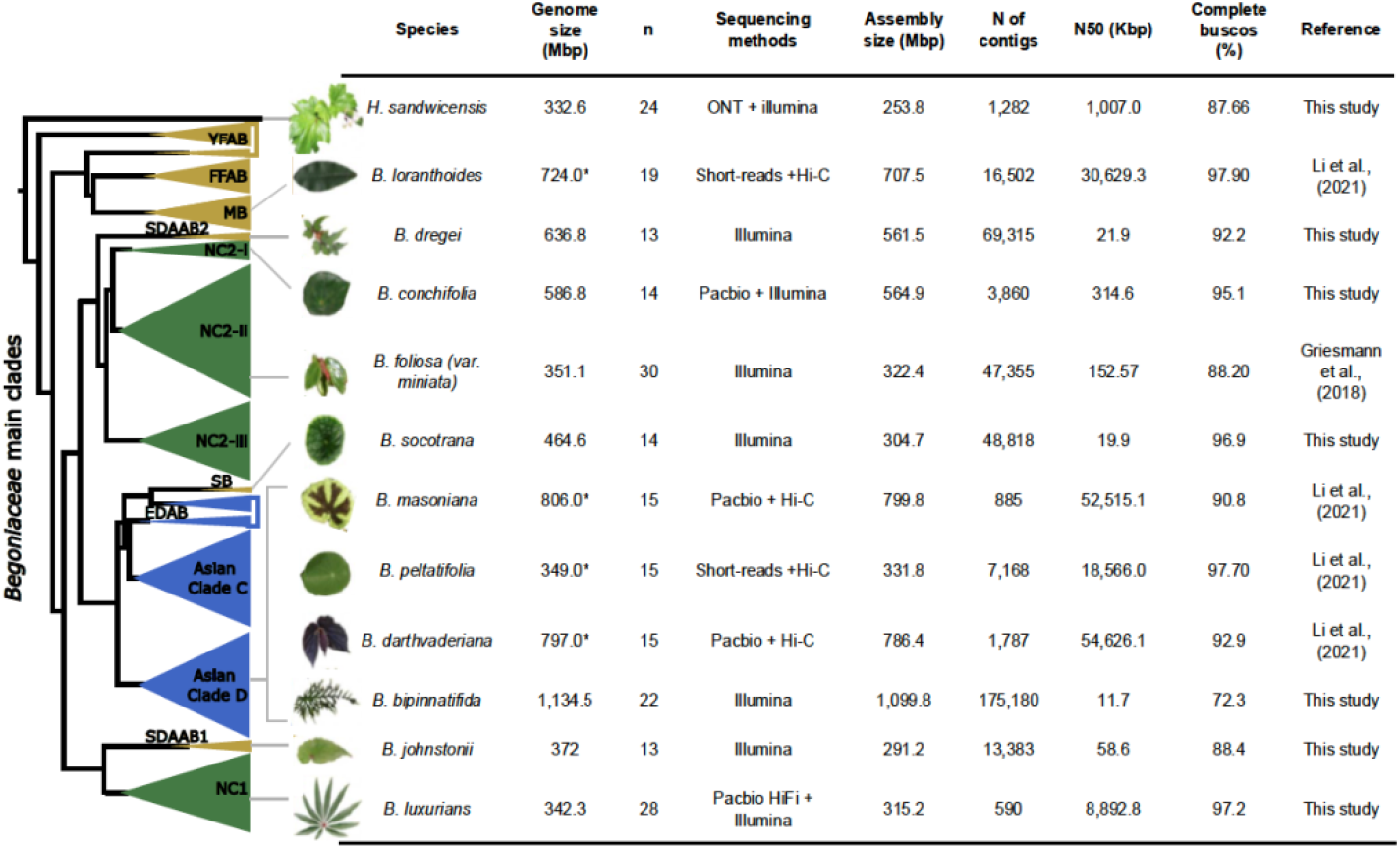
Diagrammatic representation of the Begoniaceae phylogeny (based on Moonlight et al., 2018) showing the placement of the species with sequenced genomes. All clades are supported by posterior probabilities greater than 0.85 and ML bootstrap values > 0.75, with the exception of FFAB, SB, EDAB and Asian Clade C, which have posterior clade probabilities < 0.85 and ML bootstrap values < 75. African clades are shown in yellow, Asian clades in blue and New World clades are in green. In each case, information such as genome size (1C-value) from flow cytometry data (except those marked with an asterisk (*) which come from k-mer estimations), chromosome number (n; according to Campos-Dominguez et al., 2022), and assembly statistics are given in the table.

Species assembled with long-read data (*H. sandwicensis, B. conchifolia*, and *B. luxurians*) show significantly better assembly quality statistics, similar to those for the species sequenced by Li et al. (2022). The new genomes represent a wide range of chromosome numbers (from n = 13 for the African *B. johnstonii* and *B. dregeii* to n = 28 for *B. luxurians*) and genome sizes (from the 332 Mbp of *H. sandwicensis* to the 1.13 Gbp of *B. bipinnatifida*).

### Genus-wide whole genome repeat landscapes

For each of the Begoniaceae genome assemblies, we annotated their TE content and classified them into different TE lineages (Zhang et al., 2022). Figure 2A shows the relative genome proportions of all major Class 1 and 2 TE families that comprise over 1% of the genome in at least one of the genome assemblies, as well as the total genome percentage annotated as TEs in each species (grey boxes in Figure 2a). Since the two major types of long terminal repeat (LTR) retrotransposons (i.e. Ty1/Copia and Ty3/Gypsy elements) comprised the largest proportion of the genome in most species, these were analysed in more detail by estimating the genome proportions of the different lineages LTR retrotransposons, and their genome proportions are shown in Figure 2B. Our results show variable contents for different TE families. *B. masoniana* had the largest and most TE-rich (72.65% repetitive) whereas *B. luxurians* had the smallest (28.38% repetitive) with less TEs, based on the TE classification system established by Wicker et al (2007). This, as expected, correlates to their genome size differences (except for *B. bipinnatifida*, which is likely to be a polyploid). Within the repetitive fractions, the presence of Class 1 DNA TE elements such as TIR, CACTA and Mutator, as well as some Class 2 LTR retrotransposon lineages such as Ty1/Copia-*SIRE* and Ty3/Gypsy-*Tekay* were particularly variable among species.

**Figure 2.**
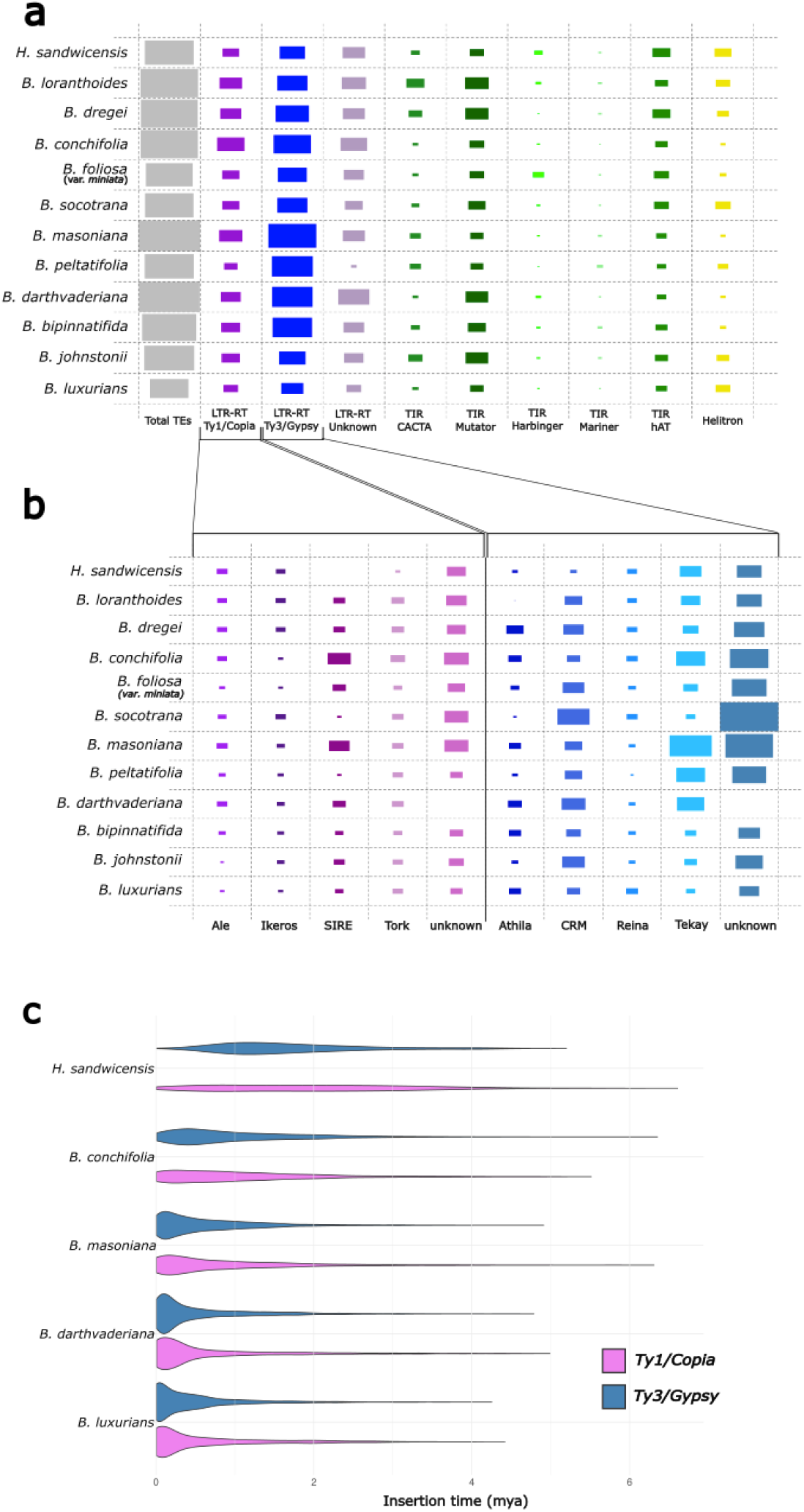
TE annotation results from analysing all available Begoniaceae genomes (sorted as in Figure 1). A. Comparative plots showing the genome percentage occupied by major TE families in each of the species analysed, arranged phyogenetically as in Fig. 1. The coloured area in each square is directly proportional to the TE content of each type. B. Comparative plots showing the genome proportion occupied by different LTR retrotransposon lineages. C. Violin plots showing the density of LTR retrotransposons inserted over time for all Begoniaceae genomes assembled with long-read data.*In-depth repeat analysis across two large Begonia species radiations*

To further explore differences in TE dynamics across species, we analysed the insertion times of intact Ty3/Gypsy and Ty1/Copia LTR retrotransposons found in each analysed genome (Figure 2C). We found that LTR retrotransposon insertions were older in the *Hillebrandia* genome compared with any *Begonia* (Figure 1c). Within *Begonia*, their intact elements seem older (2-3 MYA) in species from small groups such as *B. loranthoides, B. dregei, B. socotrana, B. peltatifolia*, and *B. johnstonii*, whereas some other *Begonia* genomes from species from large radiations showed more recent bursts of LTR retrotransposition (0-1 MYA in *B. conchifolia, B. masoniana, B. darthdvaderiana*, and *B. luxurians*). Lineage-level classification of these intact elements showed that these bursts were from different LTR lineages depending on the species. For example, in *H. sandwicensis* most of the intact LTR-RTs are Ty3/Gypsy-*Reina* elements. These are also fairly abundant in *Begonia*, but in *Begonia* other LTR retrotransposon lineages such as Ty3/Gypsy-*Tekay* and *Athila*, and several Ty/1Copia lineages such as *Ale, Ikeros, Tork, Ivana*, and *SIRE* showed specific expansions in different *Begonia* clades. Among these, Ty1/Copia-*SIRE* and *Tork*, and Ty3/Gypsy-*Tekay* and *Athila* show specific bursts in species from larger radiations (e.g., *B. conchifolia, B. masoniana, B. darthvaderiana, B. bipinnatifida*, and *B. luxurians*).

To investigate possible links between recent TE activity and *Begonia* species radiations we selected two of the largest sections of the genus: B. sect. *Gireoudia* and B. sect *Coelocentrum*, which are represented in Fig. 1 and 2 by *B. conchifolia* and *B. masoniana* respectively, and have two of the most repetitive genomes. Figure 3 shows the classification and amounts of repetitive DNA of 32 *Begonia* species from the *Gireoudia* section (Fig. 3 A) and 16 from the *Coelocentrum* section (Fig. 3 B), in their phylogenetic context. For comparison, in the case of *Gireoudia* we included two species (*B. oaxacana* and *B. heydei)* from its depauperate sister section (B. sect. *Parietoplacentalia*) comprising only three species.

**Figure 3.**
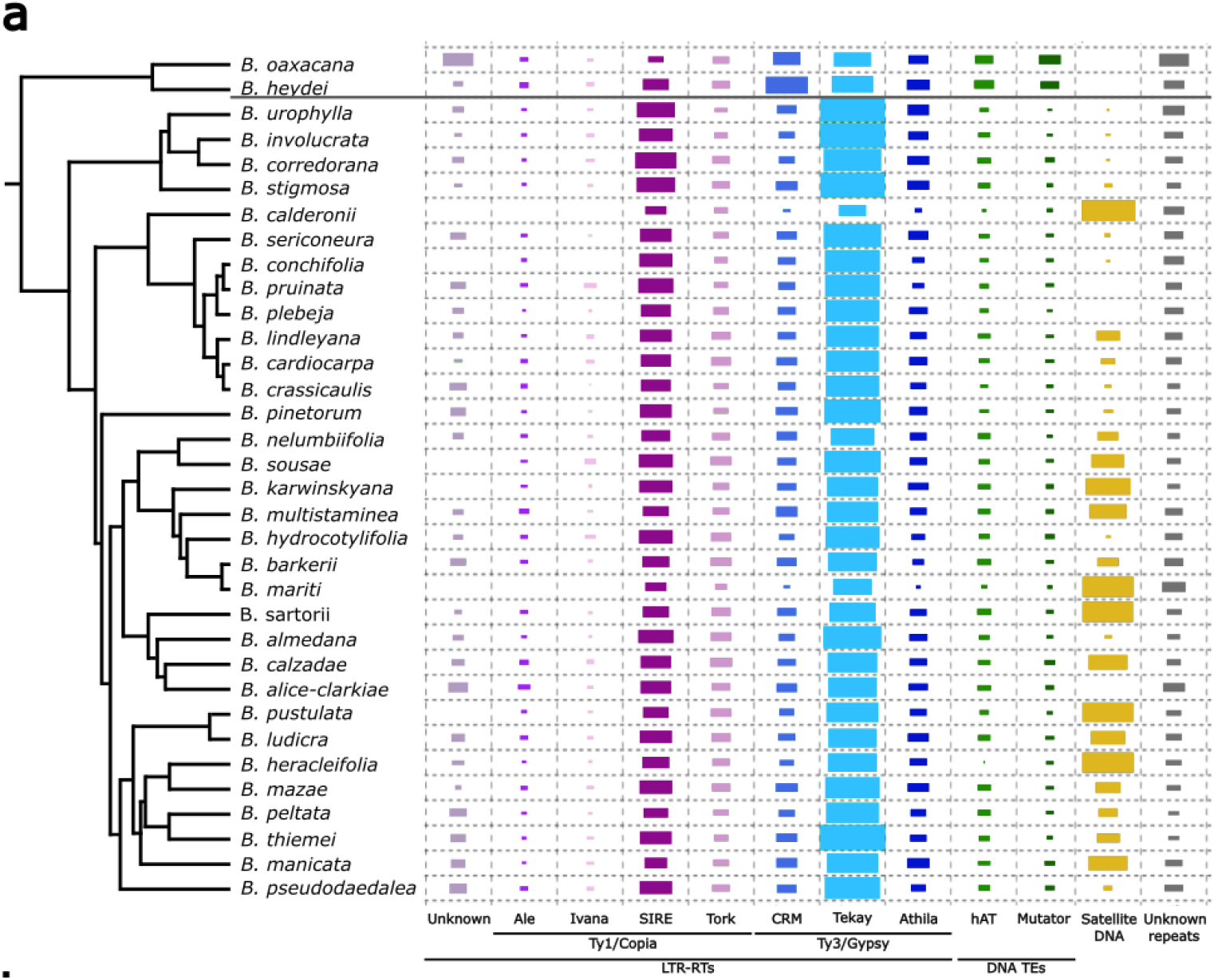
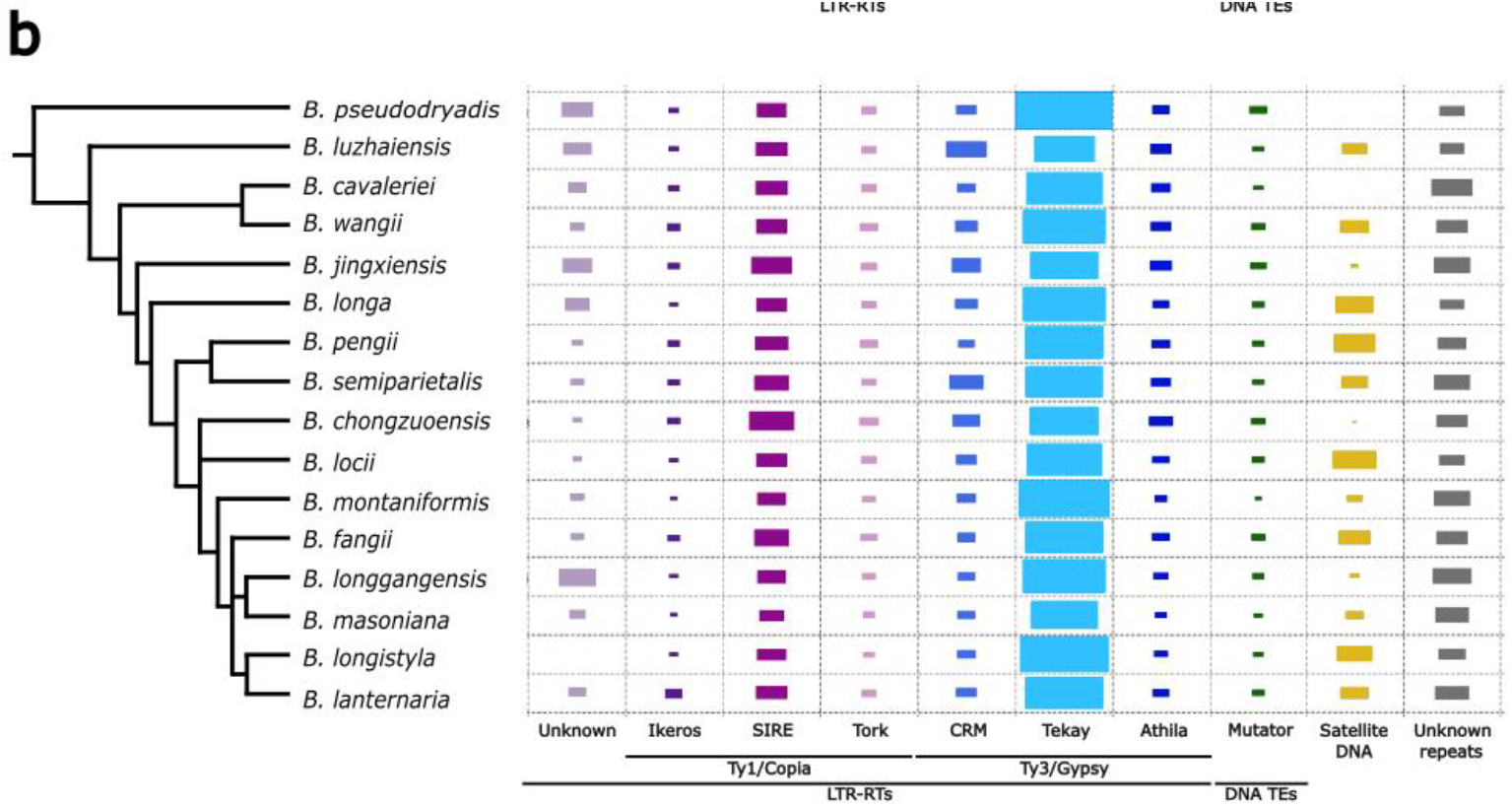
Distribution of percentages of reads clustered and annotated as different types of repeats by RepeatExplorer2 in each of the species analysed and their phylogenetic positions within the section. Phylogenetic relationships are modified from Moonlight et al. (2018). Only repeat families represented by over 1% of the reads in at least one species are shown. 2a. *Gireoudia* and *Parietoplacentalia* species (section bootstrap value >0.85). 2b. *Coelocentrum* species (section bootstrap value > 0.75).

Our repeat clustering analyses showed different Ty3/Gypsy and Ty1/Copia lineages varied in abundance (Figure 3), with most of the variable TE fraction clustered as LTR retrotransposons belonging to Ty3/Gypsy-*Tekay* (6 to 40%), and Ty1/Copia-SIRE (4 to 15%) elements, similar to the pattern seen across the genus (Figure 2). Both these families are more abundant among species from section *Gireoudia* than in species belonging to their sister section *Parietoplacentalia*. Less abundant but also comprising between approximately 0.5 and 5% of the data in most species were the LTR Ty1/Copia-*Tork*, and LTR-Ty3/Gypsy *CRM* and *Athila* families. Class 2 DNA elements were slightly more abundant in the outgroup sections (3-4%) compared to *Gireoudia* species (0-1.5%).

Figure 3 also shows that Satellite DNA (satDNA) was extremely variable across the species analysed for both sections. Percentages varied between 0% in *B. alice-clarkiae, B. pruinata* and *B. plebeja*, and 25.7% in the closely related *B. calderonii*. These values were highly variable even within *Gireoudia* sub-sections or groups, showing species-specific expansions of two clusters of satellite DNA which suggests very recent and likely ongoing expansions of satDNA families. The variation in satDNA content found corresponded to ∼8-10% of the genome size, which correlates with the genome size variation values shown for these sections in Campos-Dominguez et al. (2022) suggesting that differences in satDNA content are the main cause of genome size variation in the *Gireoudia* and *Coelocentrum* radiations.

### Intra-species repeat expansions trigger genome size variation and are associated with genetic isolation in Begonia heracleifolia

To explore the genome dynamics during incipient *Begonia* speciation, we analysed the repetitive DNA from divergent populations within the species *Begonia heracleifolia*. Previous research (Twyford et al., 2014) identified significant genetic structure and differentiation among populations in *B. heracleifolia*, and a 20% reduction in pollen viability in interpopulation F1 crosses involving a divergent population (h28) that was found to have a genome size 10% larger than the species’ mean C-value. We re-estimated the genome sizes with flow cytometry to confirm these differences, and generated low-coverage Illumina sequencing of all these populations as well as F1 progeny of h28 individuals and individuals from populations with genome sizes around the species mean (bidirectional crosses in the greenhouse between individual 397 from h28 and individual 359 from h15). Figure 4a shows the geographic locations of the sampled populations, as well as representative genome size estimates and images of leaf morphology. Figure 4b shows the repeat clustering results for the individuals analysed from across these populations. The proportions of specific LTR retrotransposon lineages and satDNA in each population were found to be variable and higher in h28 and F1 individuals compared with the other populations examined. The proportions of analysed reads clustered as satellite repeats varied from 10.6% in accession 314 (also known to have the smallest genome of the individuals analysed, as seen in Figure 4a) to 25% in accessions 397(x, greenhouse self-cross) and 400. Satellite DNA content differences were found to correlate with genome size differences, suggesting satellite DNA expansions, rather than LTR retrotransposons were mainly responsible for the difference in genome size of h28 individuals.

**Figure 4.**
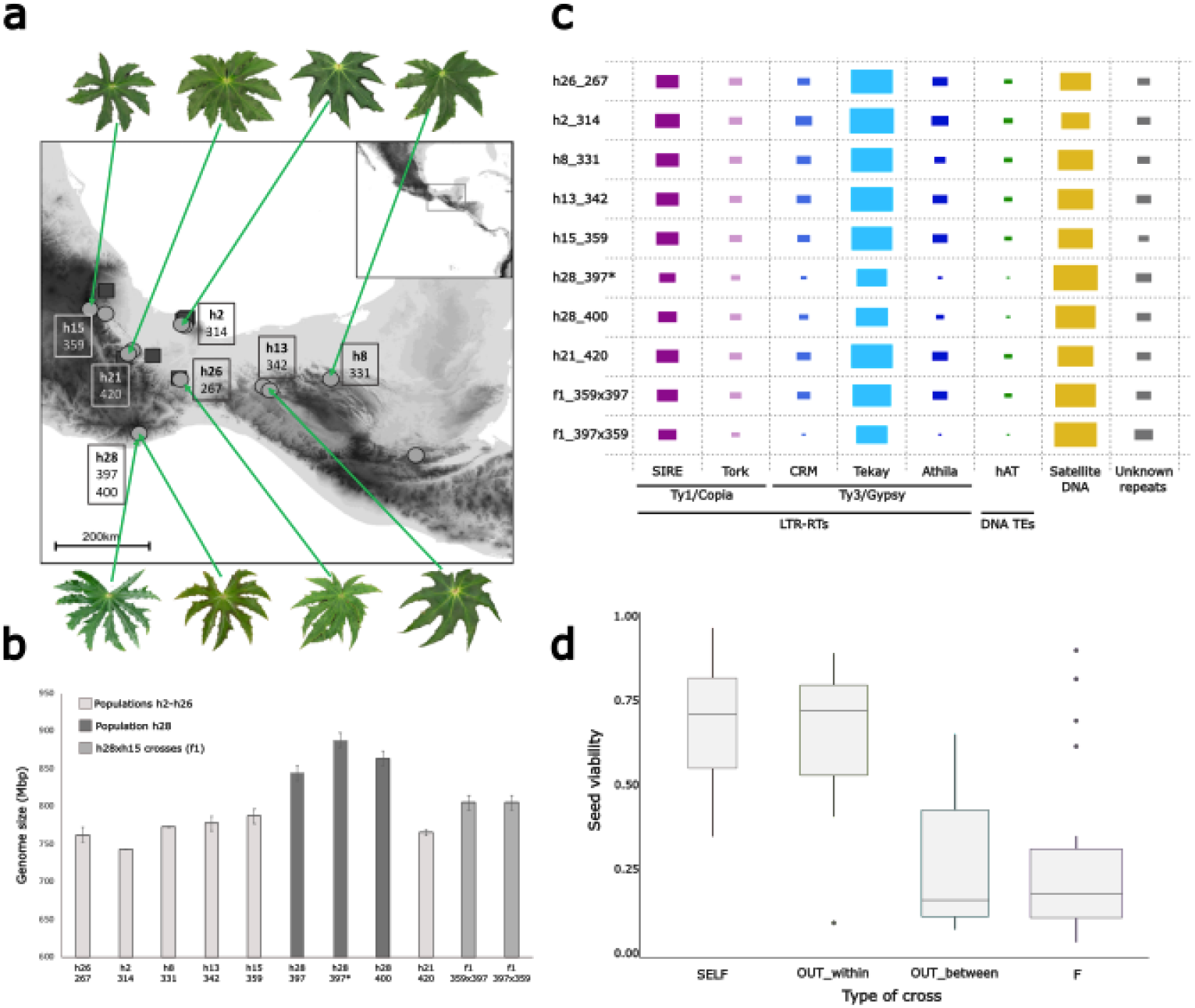
Associations between genome size, repeat content and crossing success in the widespread species *Begonia heracleifolia* in Mexico. A. *B. heracleifolia* sampling sites,genome size estimates from flow cytometry and leaf morphologies in the greenhouse. B. Repeat clustering results of the 7 populations and two F1 individuals (h28xh15 and h15xh28 greenhouse crosses, see methods). C. Box-plots representing the seed viability values of the different crosses performed between and within *B. heracleifolia* populations. Crosses are grouped along the x-axis based on the following categorisation. SELF = crosses of individuals to themselves, OUT_within = crosses of lines from within the group of populations with low genome sizes or from within the population with high genome size, OUT_between = crosses of lines from the population with high genome size with lines from the populations with low genome sizes, F = crosses involving F1s from OUT_between crosses.

We also performed RepeatExplorer2 comparative analysis between h28 (397(x)) and h15 (359) individuals. Data from these comparative analyses showed that the largest clusters with major differences between populations were satDNA and LTR-Ty3/Gypsy-*Tekay* elements. This, together with the results shown in figure 4c, indicates that not only satDNA expansions are responsible for the genome size differences found, but likely these LTRs retrotransponons have also expanded in h28. We found that these expansions were retained in the h28xh15 F1s. These F1s showed intermediate genome sizes and satDNA content compared to their two parental populations (fig 4b), suggesting Mendelian segregation patterns.

To investigate the role of these expansions in population divergence, 174 inter-and intra-population crosses were performed, and 65 capsules were sampled for seed viability counts. Figure 4c shows the distributions of the proportion of viable seeds in all analysed crosses, classified as self-crosses (SELF), crosses of lines from within the group of populations with low genome sizes or from within population h28 (OUT_within), crosses of lines from population h28 with lines from the populations with low genome sizes (OUT_between) or crosses involving h28xh15 F1s (F).

As previously observed (Twyford, 2012), self-crosses were found to be highly successful (mean proportion of viable seeds = 0.69, SEM = 0.089, n = 7). Similar crossing success was found for the OUT_within crosses (mean proportion of viable seeds = 0.66, SEM = 0.059, n = 15). The OUT_between crosses showed markedly reduced success (mean proportion seed viable = 0.25, SEM = 0.052, n = 15) Similarly, low success was seen for the crosses where one parent has a h28xh15 F1 (mean proportion seed viable = 0.26, SEM = 0.050, n = 28). The difference between the two groups of crosses with high mean seed viability and the two groups of crosses with low mean seed viability was supported by non-parametric analysis of variance (Kruskal-Wallis test, SELF and OUT_within vs. OUT_between and F, p < 0.0021). Overall, this suggests that population h28 exhibits reduced compatibility with other populations of *B. heracleifolia* and that it may be in the early stages of reproductive isolation.

## Discussion

In this study we explore the role of a dynamic genome in species radiations in the genus *Begonia*, which is exceptionally species-rich and has several parallel radiations in Asia and the Americas. These particular characteristics have allowed us to look across the genus for correlations between repeat dynamics and species radiations. We focus in on the species *B. heracleifolia* which has a wide distribution and a large range in genome size between individuals, making it a tractable experimental system to examine in detail genomic events accompanying a putative on-going speciation event. Three of the genomes presented here are new high-quality assemblies based on long-read data, including the sister species to *Begonia* (*H. sandwicensis*) as well as two representatives from Neotropical *Begonia* clades (*B. conchifolia* and *B. luxurians*). Our analyses of the genomic content of repetitive DNA in a few (12 with whole genome assemblies + 50 with low-coverage data) taxonomically representative *Begonia* species from independent radiations provide a powerful overview of the repeat landscape across a wide phylogenetic range of species, including two large radiations and one diverging species (*B. heracleifolia*; Twyford et al., 2014).

Our genomic analysis of *Hillebrandia sandwicensis* reveals that this species has a smaller, less repetitive and more stable genome (with less recent LTR activity) than all *Begonia* species analysed. Clearly these sister lineages have experienced extremely different fates since they diverged, which may be as long as >55 million years ago (Forrest, 2001) or may have occurred more recently (25 MYA, Moonlight et al. 2018). The high homozygosity (observable in the k-mer profiles in supp. fig X) of this wild collected accession suggests populations of *Hillebrandia* in its native range are small and inbred. *Hillebrandia* was categorised as in peril of extinction in 1999 (according to NatureServe), supporting the conclusion that it is suffering from severe inbreeding depression, with this taxon potentially at an ‘evolutionary dead end’ (Lande Schemske, 1985). If, as suggested by Cronk (1989), taxa go through ‘boom and bust’ cycles, *Hillebrandia* may be the last dying member of what once was a very species-diverse plant group (Hughes, 2002). Further genetic analyses and additional population structure studies on *Hillebrandia* would be needed to confirm its declining evolutionary “success”.

Within *Begonia*, we found different genome dynamics in each of the radiations studied, but with one clear pattern: species from large radiations show an expansion of at least one repeat family compared with their close relatives which did not experience an extensive radiation. From the first Neotropical radiation (NC1, Figure 1), we sequenced and analysed the genome of *Begonia luxurians*, from the large section *Pritzelia*. Species from this section, including *B. luxurians*, have a large number of chromosomes (2n=30-70, with 2n=56 in ∼60% of the species with known data for this section; Campos-Dominguez et al., 2022), and the smallest genome sizes found so far reported in *Begonia* (245 to 345 Mbp/1C), some even smaller than *Hillebrandia* (DeWitte et al., 2009; Campo-Dominguez et al., 2022). The high chromosome numbers in all sections from this clade suggest a polyploidisation event early in the diversification of this group. However, the small genome sizes indicate extensive genome downsizing after such an event. Wang et al., (2021) describe that this phenomenon has been observed in many polyploid species, and suggests this could be caused by various recombination pathways and leading to loss of different types of DNA sequences (e.g. repeats, genes, low-copy, non-coding DNA). However, although polyploidisation events have been suggested for other *Begonia* clades (e.g. NC2-ii in Campos-Dominguez et al.,2022), other sections do not show the levels of genome downsizing seen in *Pritzelia*. This suggests either very early genome downsizing directly following polyploidisation in the common ancestor of this section, or some selective pressure for small genomes for these species. Wang et al. (2021) and Leitch and Bennett (2004) also discuss that DNA loss after polyploidy could also be under environmental selection driven by the need to reduce nutrient costs (N and P) associated with DNA synthesis, as well as the costs of building and maintaining the increased cell size of a polyploid. Obtaining additional genome size data and further genomic and ecological analysis on *Pritzelia* species would provide useful information for understanding genome size evolution in this section. The genome size variation observed among *Pritzelia* species, our repeatome annotation and LTR dating results suggest that although their LTR content is low, these species show greater genome dynamism compared to their closest African relative (*B. johnstonii*) in the species-poor section *Rostrobegonia*.

Regarding the second Neotropical radiation (NC2), we analysed the complete genomes of two species: *B. conchifolia* (NC2-i) and *B. foliosa* var. miniata (NC2-ii). Our own *B. foliosa* data and data from Campos-Dominguez et al. (2022) suggest potential polyploidisation events in NC-ii, and although genome sizes are not as small in NC2-ii as in NC1, they are smaller than those in sister clades and have higher and extremely variable chromosome numbers (2n=22-156, whereas NC2-i and NC2-iii show 2n=28 in most of their species with known chromosome numbers). *B. conchifolia* from the NC2-i radiation, on the other hand, does have greater LTR retrotransposon content and a slightly larger genome (the largest of the three Neotropical genomes analysed) than both other Neotropical species and its depauperate sister *B. dregeii*, and this could be due to higher LTR retrotransposon activity. Our additional results from section *Gireoudia* species and *B. heracleifolia* populations (see below) suggest that satDNA dynamics are also important in this radiation.

Regarding the Asian *Begonia* groups, our TE annotations using full genomes from species in sections *Coelocentrum* (*B. masoniana*) and *Petermannia* (*B. darthvaderiana* and *B. bippinnatifida*) are representatives from two of the main radiations within the Asian clade. Campos-Dominguez et al. (2022) showed species in these sections had relatively stable karyotypes but variable genome sizes, suggesting the dynamic nature of these genomes. Here, TE annotation results on the genome assemblies indicate high content and recent activity of LTR retroelements in *B. masoniana, B. darthvaderiana* and *B. bipinnatifida*, in contrast with the repeat landscape of *B. peltatifolia*, which has a considerably smaller genome. Although there is currently no phylogenetic information (Moonlight et al., 2018), nor population data published for this species, *B. peltatifolia* is mostly found in small, isolated populations distributed across a compact range (Xiaobai (1993) and M. Hughes, pers. observ.). Moreover, this species is categorised as endangered according to the IUCN (International Union for the Conservation of Nature, 2017; Tian et al. (2018)). It could be possible that within *Begonia*, there are similar cases to that of *H. sandwicensis*; i.e. highly isolated species with stable genomes in an “evolutionary dead end”. This suggests that the factor(s) responsible for a dynamic genome in large *Begonia* radiations are not present in all *Begonia* species.

Our genomic analyses of the two large *Begonia* species radiations provide a better overview of genome dynamics at a closer phylogenetic level. These are both large monophyletic clades with over 90 species each and stable chromosome numbers, but variable genome sizes (Campos-Dominguez et al., 2022). Our results indicate that the genome size variability found in *Gireoudia* and *Coelocentrum* species is caused by different LTR-Ty3/Gypsy, LTR-Ty1/Copia and satDNA expansions. These results contrast with the repeat landscape of the smaller, closely related clades, which have less repetitive genomes (especially in terms of the abundance of retrotransposons and satDNA). This provides correlative evidence that genome dynamism may play a role in species radiation in larger *Begonia* sections. In addition, our study of *B. heracleifolia* populations with different genome sizes indicates that the large intra-specific genome size differences here are driven predominantly by satDNA and LTR-Ty3/Gypsy-*Tekay* dynamics. Interpopulation crosses between larger and smaller genome size individuals within this species yielded fewer viable seeds, indicating a potential role of these repeats in reproductive isolation and speciation.

Our findings suggest a link between large radiations and higher rates of repetitive DNA gain/loss, but we acknowledge it is difficult to identify the specific factors directly triggering speciation or diversification events when limited to studying a few genomes from radiations of hundreds of species. Previous research has linked diversification and TE activity in other taxa, such as for example African cichlid fishes (Carleton et al., 2020), *Hydra* species (Wong et al., 2019) and recently cultivated olive tree varieties (Jimenez-Ruiz et al., 2020). However, the approaches taken were different in each case, and are therefore not applicable to the study of such a large genus with so many different species radiations. Moonlight (2017) studied the link between *Begonia* speciation and niche variation, and his results revealed that the studied Neotropical clades did not follow adaptive radiations. Instead, non-ecological speciation was suggested for some *Begonia* groups such as the *Gireoudia* radiation. This evolutionary model aligns with our results, which also suggest genetic drift and sudden structural genomic changes contribute to speciation and shape species radiations in *Begonia*. Nonetheless, this genus has undergone several radiations, and more in-depth evolutionary studies are needed in each of them to fully understand the role of different ecological and genomic factors in each event. Naciri and Linder (2020) reviewed the different scenarios and factors that can lead to different adaptive radiations, including the arrival to a new adaptive zone, geographical and ecological expansion, epigenetic changes, genetic drift, gene flow limitation, admixture or hybridisation, and all of these could interact with or drive the activation and dynamics of repetitive DNA. Evolutionary radiations are complex processes and although TEs represent an important source of genomic variation, it is essential to consider the evolution of *Begonia* in a broad context in which numerous evolutionary scenarios could have triggered the many radiations represented by its largest sections. Nevertheless, we suggest the patterns we report here support a role for genome dynamics and TE activity in Begonia radiations.

Satellite DNA expansions across populations have also been described in other species. Shah et al. (2020) found remarkable satellite DNA expansions among populations of grasshoppers, and suggested that such expansions were not the cause of genome size expansions but rather the consequences of selection as a means of stabilising the genome previously expanded by LTR retroelements. This theory relies on other genomic factors triggering the genome expansion, and in the case of *B. heracleifolia* such factors could include the LTR-Ty3/Gypsy-*Tekay* lineages which we showed expanded in h28 (Fig. 4). Another potential scenario could be that the satellite DNA expansion was triggered by transposition. Until recently, there was no evidence of the functional and structural links between transposons and satellite DNA (Garrido-Ramos, 2017). However, in recent years research suggests that TEs can act as substrates for origin and mobility of satellite DNAs (Dias et al., 2014; Mestrovic et al., 2015; McGurk and Barbash, 2018), linking the spread of short tandem repeats to transposition processes (Satovic et al., 2016). This link between two key factors in genomic change provides a new perspective on genome evolution that suggests transposon insertion and recombination dynamics can contribute to satellite DNA formation and variation. Henikoff et al (2001) proposed a “centromere paradox”, a theory on why we might expect centromeric repeats to be fast-evolving and cause cross incompatibilities between species. Rodriguez et al (2017) attributed a possible role in speciation to these elements because of the species-specificity of the satDNA families (Rodriguez et al., 2017), and another recent study (Schmidt et al., 2024) also found a direct association between genome size and species-specific satellite DNA dynamics in beets (*Beta* and *Patellifolia*). A more in-depth satellitome analysis of these species and the *B. heracleifolia* populations (e.g. as done for *Vaccinium* in Sultana et al. (2020) or for the ladybird beetle in Mora et al. (2020)) would help us confirm whether these satDNA differences can be attributed to centromeric regions, which would support the idea of centromere divergence triggering genomic incompatibilities and problems in chromosome segregation as suggested in Henikoff et al (2001).

Speciation is a highly complex process affected by numerous factors. Studying a continuous process brings some challenges and leads to many open questions. The main question after considering all our results would be whether the described genomic changes are sufficient to drive the partial reproductive isolation we found towards a stronger isolation. Kulmuni et al. (2020) discuss that this transition is driven by many factors that could be either ecological or of any other nature, but leaves a detectable signal in the genome. Serrato-Capuchina and Matute (2018) discuss the role of TEs in reproductive isolation and provide examples that mostly rely on adaptive phenotypic changes, and conclude that there is still little direct evidence that TEs can facilitate reproductive isolation and ultimately speciation. To determine whether DNA repeats and TE dynamics actually drive speciation in *Begonia*, are a response to these processes, or are merely neutrally associated, further studies are needed. These should include higher quality genomes across the genus and population level analysis of both genomic changes and gene flow, along with a deeper understanding of the biotic and abiotic factors that might shape patterns of genetic diversity in these species.

## Acknowledgements

The authors would like to thank Mike DeMotta and Dustin Wolkis from the NTBG for sharing the *Hillebrandia* plant material people, and Alex Monro (RBGK) for sharing *B. oaxacana* dry material. Thanks also to Andrés Romanowski for allowing us the ONT sequencing of B. conchifolia, which greatly contributed to improve the assembly. Thanks to the RBGE horticultural staff, for mantaining the Begonia living collection. We also thank the laboratory team at AllGenetics & Biology SL, who performed the DNA extraction, and library prep and sequencing of *B. oaxacana, B. heydei* and *B. pinetorum*. The authors acknowledge the Research/Scientific Computing teams at The James Hutton Institute and NIAB for providing computational resources and technical support for the “UK’s Crop Diversity Bioinformatics HPC (Percival-Alwyn et al., 2024)”, (BBSRC grant BB/S019669/1), use of which has contributed to the results reported within this paper.

## Author contributions

LCD and CAK designed and conceived the project, and wrote the manuscript. LCD performed or supervised all the assemblies and bioinformatic analyses in this study. TEK conducted the DNA extractions, sequencing data analysis and genetic crosses of the *B. heracleifolia* populations. LC performed the analysis of the *Coelocentrum* species data. AMM and MD generated the *Hillebrandia* genome assembly. CF did the high-molecular weight DNA extractions for *B. conchifolia* nanopore and *B. luxurians* Pacbio sequencing. Y-HT, HAQ, L-ND and K-FC generated most of the *Coelocentrum* and *Gireoudia* sequencing data. ABG contributed to the *B. conchifolia* sequencing and all genome assemblies, as well as to the project planning and discussions. JP performed the *B. heracleifolia* genome size measurements. JP, IL, ABG and ADT contributed to manuscript writing and project discussions.

## Data availability

All scripts written for the genome analysis are kept in the following repository: https://github.com/Lcamdom/Begonia_genome_analysis. The raw genomic data generated for this study are stored in [
SRA code]. Genome assemblies are stored [here].

